# Dissecting Key Multivalent Processes in Glycosidase Inhibition: Insights from Thermodynamic Modelling and Atomistic Simulations

**DOI:** 10.1101/2023.11.29.569169

**Authors:** Martin Spichty, Yan Liang, Rosaria Schettini, Irene Izzo, Anne Bodlenner, Philippe Compain

**Affiliations:** Laboratoire d’Innovation Moléculaire et Applications (LIMA), University of Strasbourg | University of Haute-Alsace | CNRS (UMR 7042); Equipe Chimie Théorique et Modélisation Biomoléculaire (CTMB); IRJBD, 3 bis rue Alfred Werner, 68057 Mulhouse Cedex, France; Laboratoire d’Innovation Moléculaire et Applications (LIMA), University of Strasbourg | University of Haute-Alsace | CNRS (UMR 7042), Equipe de Synthèse Organique et Molécules Bioactives (SYBIO), ECPM, 25 Rue Becquerel, 67087 Strasbourg, France; Department of Chemistry and Biology “A. Zambelli”, University of Salerno, Via Giovanni Paolo II,132, 84084 Fisciano, Salerno, Italy

**Keywords:** multivalency, multivalent inhibitory effect, thermodynamic modelling, molecular dynamics

## Abstract

Multivalency represents a powerful approach to increase the inhibition potency of moderate glycosidase inhibitors. Regarding the key role of catalytic glycoside hydrolysis in biology, understanding the molecular mechanisms and origin of the multivalent inhibitory effect is of great interest and presents a fascinating playground for theoretical studies. Our teams have recently dissected key processes of multivalent glycosidase inhibition through the use of different neoglycoclusters based on deoxynojirimycin (DNJ) inhitopes and a cyclopeptoid scaffold. This companion article details the theoretical aspects of this former study. A thermodynamic model is developed and validated, compared to literature, and extended to account for particularities of the charged DNJ inhitopes.

## Introduction

Glycosidase inhibitors have found applications as agrochemicals and therapeutic agents, for example, to target viral infection, cancer, and genetic disorders.^1,2^ A promising strategy to increase the potency of such inhibitors is the use of multivalent clusters, molecules that cluster together multiple substrate mimicking molecular units (inhitopes) that can bind to and thereby inhibit the corresponding glycosidase enzyme.^3,4^ The neoglycocluster **1** represents one of the most potent examples of this kind. The number of inhitopes per cluster (= valency *n*) is an important parameter for the inhibitory effect. Thereby the inhibition can increase in a non-linear fashion with increasing *n, i*.*e*., the increase of the valency by an order of magnitude can increase the inhibition potency by several orders of magnitude. For example, the iminosugar 1-deoxynojirimycin (DNJ) and related *N*-alkylated derivatives such as **2** are relatively modest inhibitors^4^ of the enzyme Jack Bean α-mannosidase (JBα-man): the divalent cluster **7** (Scheme 1) with two DNJ units displays an inhibition constant of 54 μM (Table 1). ^5,6^ The 36-valent cluster **1**, however, features an inhibition constant of only 1.1 nM.^7^

**Table 1:** Experimental^6^ and theoretical data for the studied multivalent clusters.

| Cluster | Valency $n^a$ | $\Omega = n(n-1)^b$ | $K_i$ (nM)<sup>c</sup> | $rp$ (exp)<sup>d</sup> | $rp$ (theor)<sup>e</sup> |
| --- | --- | --- | --- | --- | --- |
| **1** | 36 | 1188 <sup>f</sup> | 1.1 | 49091 | 45362 |
| **3** | 12 | 132 | 74.0 | 730 | 970 |
| **4** | 12 | 132 | 42.0 | 1286 | 970 |
| **5a** | 12 | 108 <sup>g</sup> | 84.0 | 643 | 794 |
| **5b** | 12 | 108 <sup>g</sup> | 120.0 | 450 | 794 |
| **5c** | 12 | 108 <sup>g</sup> | 97.0 | 557 | 794 |
| **6a** | 4 | 12 | 3100.0 | 17 | 17 |
| **6b** | 4 | 12 | 2200.0 | 25 | 17 |
| **6c** | 4 | 12 | 2300.0 | 23 | 17 |
| **7** | 2 | 2 | 54000.0 | $\equiv 1$ | $\equiv 1$ |

<sup>a</sup>Number of inhitopes. <sup>b</sup>Degeneracy of complex (see text). <sup>c</sup>Experimentally measured inhibition constants. <sup>d</sup>Relative inhibition potency= $K_i$ (divalent reference **7**)/ $K_i$ (cluster). Note that the companion study<sup>6</sup> uses **2** as a monovalent reference. <sup>e</sup>Calculated with Eq. 10 with $p=1.5$ and the given degeneracy coefficients. <sup>f</sup> Reduced degeneracy: $\Omega=36 \times 33$ (see text). <sup>g</sup> Reduced degeneracy: $\Omega=12 \times 9$ .

**Scheme 1.**
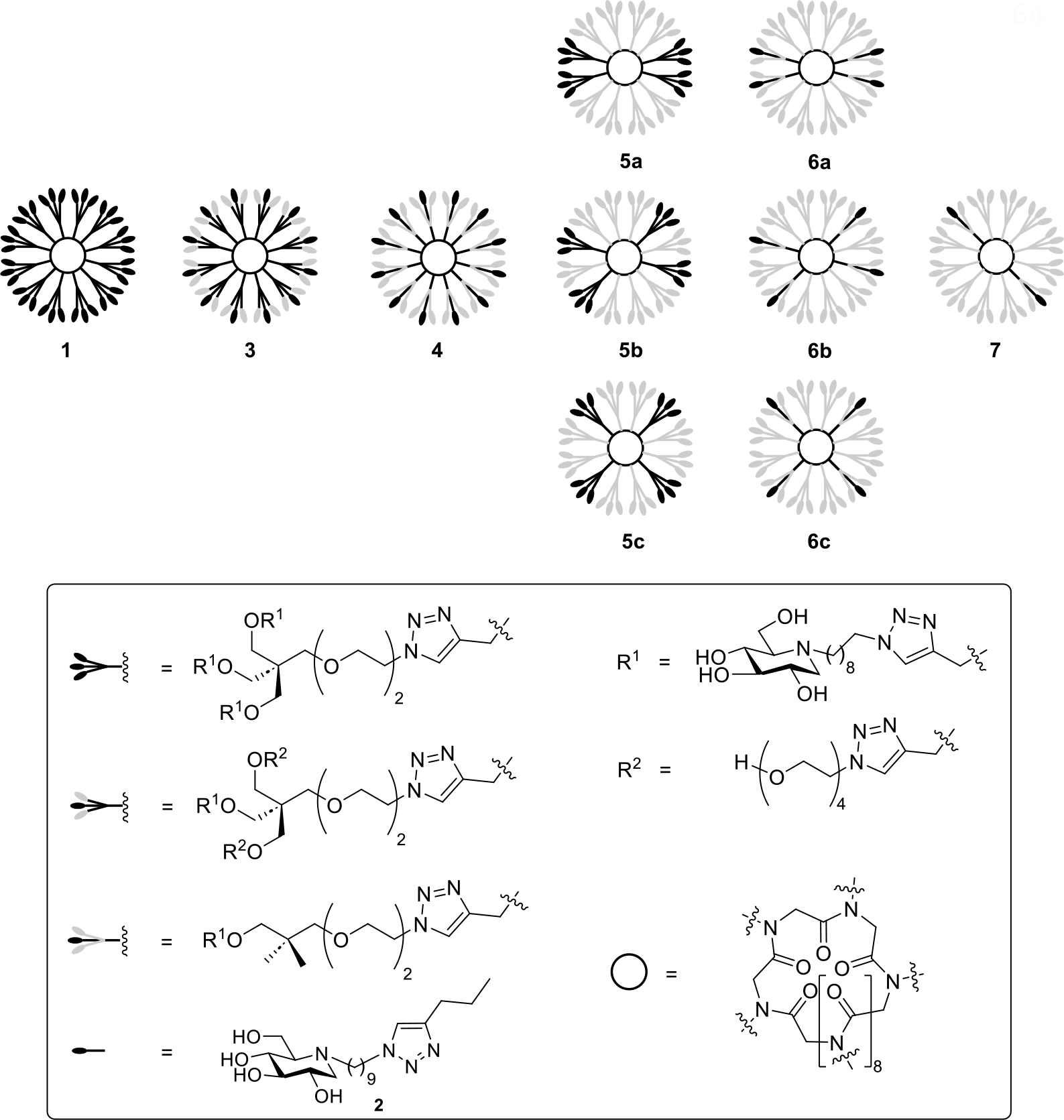
Simplified representation of DNJ-based cluster **1** and of its deconstructed analogues **3**-**7**. The molecular entities that are removed as part of the deconstruction process are shown in light grey.

Thus, the increase of the valency by a factor of 18 increases the potency by a factor of 50 000! This over-amplification of the inhibitory effect (beyond that would be expected from an analogous concentration increase) is referred to as the cluster or multivalent effect. The molecular sources of the multivalent effect are not yet fully understood. Recently, we studied by a deconstruction approach a series of new DNJ-based clusters (**3**-**7**) to investigate the architectural parameters that influence the multivalent effect when targeting JBα-man.^6^ These clusters consist of a cyclic peptide backbone where multiple branches are attached to the backbone nitrogen atoms. Each branch comprises a flexible linker and a terminal DNJ inhitope. Under experimental conditions of the inhibition measurements (pH=5) the N-alkylated DNJ is mostly protonated (pKa = 6.7-7.1).^8,9^ Clusters **3**-**7** differ not only by the number of branches but also by the orientation of the branches and the chemical structures of the linkers (*unipod* or *tripod*).^4^

The enzyme JBα-man (220 kDa) is a homodimer (LH)^2^ bearing two active sites. Its crystal structure^10^ in complex with the 36-valent cluster **1** shows that the enzyme can also form dimers with four active sites; Fig. 1A shows a schematic illustration of this dimer. Each site is bound by a DNJ unit. The formation of dimers has been confirmed by analytical ultracentrifugation sedimentation velocity (AUC-SV) experiments but only for high-valency clusters **1** and **4**.^6^ It is, however, not clear if the formation of dimers is of relevance for the measurements of the inhibition potency (since these experiments are carried out with a 1000x lower (1 μg/mL) concentration than the AUC-SV experiments (1 mg/mL).^6^ Obviously, the inhibition potency of these clusters depends strongly on the valency as seen when cluster **1** is deconstructed step by step (Table 1). But there is also an influence on the chemical structure of the branches, or more precisely of the linkers, as seen from the series with *n*=12: clusters **5a**-**c** with four *tripod* linkers feature about 1.5 times lower inhibition potencies than cluster **3** with twelve *tripod* linkers but only with one inhitope per branch. And the potency of cluster **3** is about 1.5 lower than that of cluster **4** with twelve *unipod* linkers. The importance of the linkers on the multivalent effect has been previously reported.^11^ On the other hand, the orientation of the branches has little influence (**6a**-**c**). To complement this experimental work, we briefly highlighted in the companion study some theoretical and simulation results.^6^ The aim of this article is to provide a greater in depth analysis of these theoretical and computational aspects.

**Figure 1:**
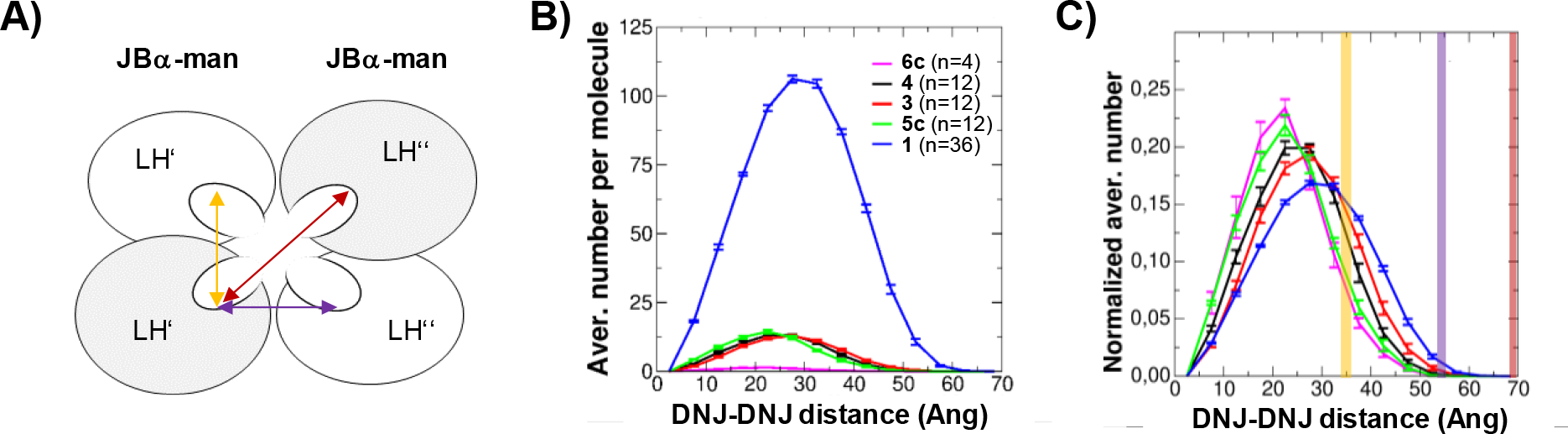
Geometric parameters of the studied system. B) Schematic representation of dimeric JBα-man as seen in the crystal structure.^10^ The distances between the active sites are indicated by arrows; their values are given in subfigure C) as vertical bars (with the same color code). B) Distribution of DNJ-DNJ distances as obtained by atomistic simulations. The positions of the nitrogen atoms of the DNJ heads were used for the calculation. C) Same as A) but normalized by *n*(*n*-1).

Thermodynamic models can help to understand the basis of multivalency effects.^12–14^ Here we develop such a simplistic model with the aid of macroscopic rate constants and equilibrium constants. We only consider the formation of complexes with 1:1 stoichiometry and we assume identical chemical activity of unbound DNJ inhitopes at all steps along the binding pathway. We discuss the relationship of our approach with the more advanced approach of Kitov and Bundle.^12^ We validate assumptions of the thermodynamic model using atomistic simulations than can provide valuable structural insights into the glycoclusters.^15,16^ We point out limitations of such thermodynamic models for the studied inhibitors, particularly, in the context of JBα-man dimerization (2:1 stoichiometry). We then apply an extension to account for the increasing net charge of the inhibitor with increasing valency.

## Results and Discussion

### A thermodynamic non-cooperative model

We start with a model for the inhibition of JBα-man protein (**P**, see Scheme 2) with two binding sites by a *n*-valent DNJ-based inhibitor (**I**) . We imply that the protein **P** and inhibitor **I** form first an encounter complex **PI** before any DNJ head binds to the active site of the protein (encounter association constant *K*_*1*_). We further imply *k*_on_ is the theoretical rate constant of DNJ inhitope binding for a monovalent inhibitor associated to a hypothetical enzyme with only one binding site; *k*_off_ is the corresponding rate constant for the unbinding of the DNJ inhitope. We also assume that the binding of the first DNJ inhitope, to form a singly occupied **PI**^**1**^ complex, does not influence the binding rate of the second DNJ inhitope to form the double occupied complex **PI**^**2**^ (no positive or negative cooperativity). As a result, the macroscopic rate constants for binding the first and second inhitopes are 2*nk*_on_ and (*n*-1)*k*_on_, respectively. The unbinding rate constants are 2*k*_off_ (1^st^ unbinding) and *k*_off_ (2^nd^). Note that in this model with macroscopic rate constants the species **PI**^**1**^ and **PI**^**2**^ comprise all possible ways to form a single and double occupied complex with a *n*-valent inhibitor.

**Scheme 2.**
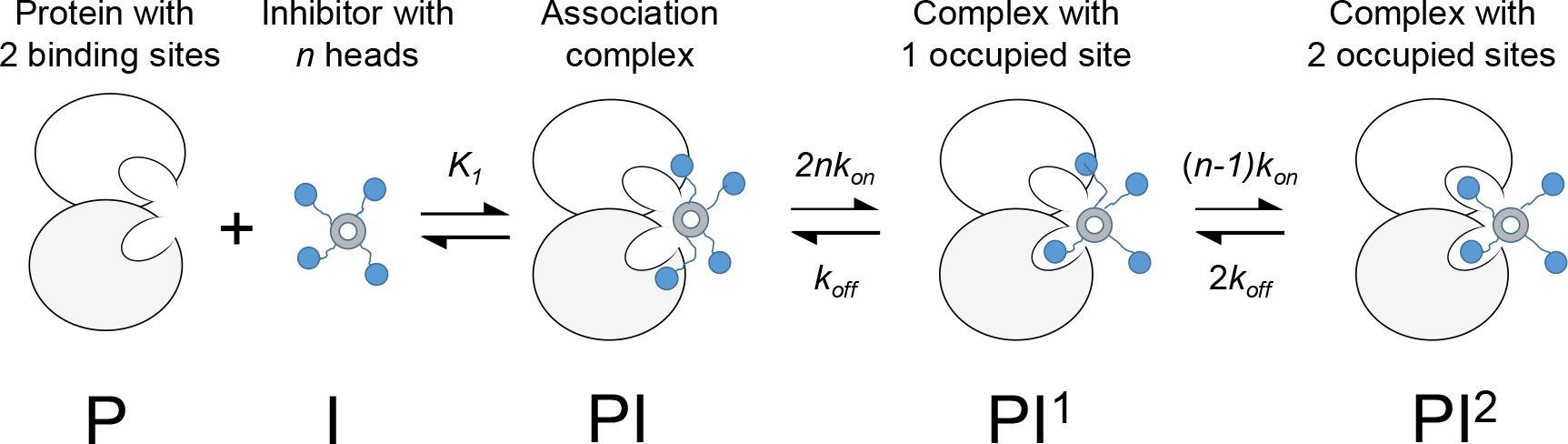

Thus, at equilibrium we find the following expressions:

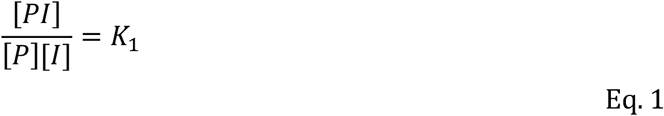

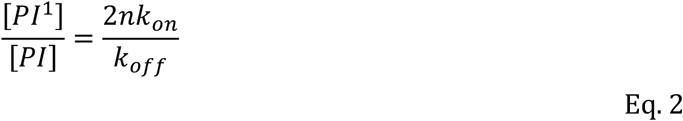

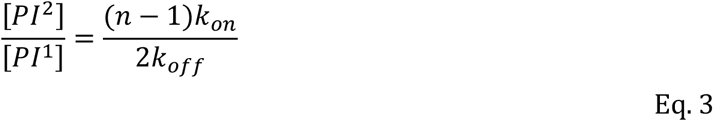

The fraction of occupied binding sites is given by:

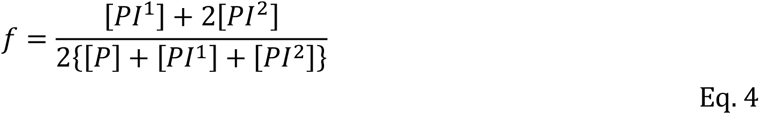

Under the condition *f*=0.5 (mimicking the experimental *K*_i_ measurement) we obtain:

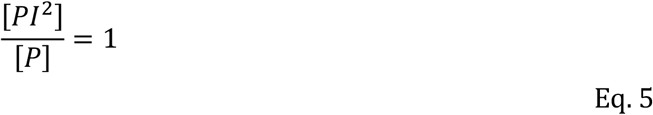

or equivalently

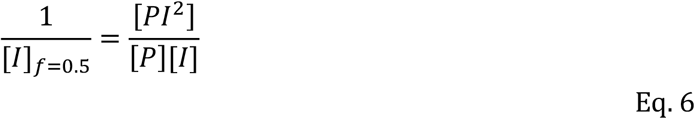

We note that the right-side is the definition of *K*_eq,*PI2*_, *i*.*e*., the equilibrium constant for the formation of the fully occupied protein (bound by two DNJ heads) and we obtain:

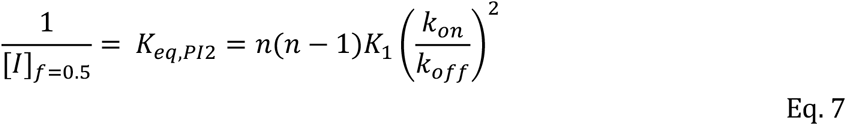

and for the relative potency *rp* we find:

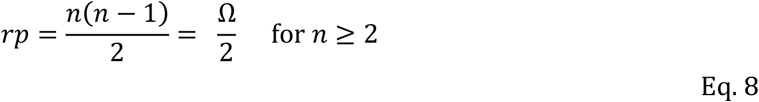

when we use the case *n*=2 with *K*_*eq,PI2*_ = 2*K*_*1*_*(k*_*on*_*/k*_*off*_)^2^ as reference.

Kitov and Bundle determined thermodynamic models for the inhibition of multisite receptor proteins by multivalent ligands.^12^ These models were developed for 1:1 stoichiometry and under the assumption that the *m* binding sites of the receptor protein and the *n* branches of the inhibitor act independently and have identical binding properties. (Note that in their publication, Kitov and Bundle used the parameters *m* and *n* interchanged.) These assumptions are implicitly included in our model, too. Kitov & Bundle developed their models for different types of topologies. Our results above can be derived from the thermodynamic models of Kitov & Bundle assuming radial topology. Radial topology implies that the *n* branches are centrally anchored so that each ligand can interact with each binding site with the same binding strength. Again, radial topology was implicitly included in our model. Under these assumptions, Kitov & Bundle (see above) showed that for a protein with two binding sites (as in the case of JBα-man, *m*=2) the equilibrium constant for the formation of fully inhibited enzyme increases with *n*(*n*-1). This *n*(*n*-1) dependence results from the **degeneracy coefficient** Ω that accounts for the fact that complex designated as **PI**^**2**^ are “not individual molecules but an ensemble of Ω microscopically distinguishable complexes”.^12^ In our model this degeneracy was incorporated into the macroscopic rate constants.

There is an illustrative approach to rationalize the dependence of *K*_eq,*PI2*_ (or equivalently of *rp*) on *n*(*n*-1) when implying radial topology. This approach offers also a way to verify the assumption of radial topology. To do so, we rely on the fact that the free energy is a state function and the free energy of binding (or *K*_eq,PI2_) does not depend on the chosen pathway. Let us consider the conformational selection pathway: two DNJ heads need to have the correct distance (and orientation, accessibility, etc) to be selected by the enzyme for binding to its active sites. According to the Gauss summation formula, the probability of finding such two DNJ heads is proportional to *n*(*n*-1)/2 if all heads are equivalent and independent from each other.

### Validation of radial topology

The branches of the studied clusters in this work are not exactly centrally anchored and the type of branches are not all identical. To probe the assumption of radial topology, we investigated the geometric properties for a subset of different clusters with the aid of full-atom molecular dynamics. We therefore determined the distribution of DNJ-DNJ distances for clusters **6c** (*n*=4), **3, 4, 5c** (*n*=12) and **1** (*n*=36). To be in line with the required assumptions of radial topology, the same protonated state of all DNJ heads was used (which can be considered as a low pH case). The raw distributions nicely visualize the *n*(*n*-1) multivalency effect (Fig. 1B): the average number of DNJ heads at the same distance as in the protein is by factors higher for *n*=36 than for *n*=12 and *n*=4.

When normalized by *n*(*n*-1) the distributions of DNJ-DNJ distances are much more similar (Fig. 1C). The distributions become, however, slightly broader with increasing valency. As a result, the normalized average number of DNJ heads at a distance of ca. 35 A (= separation of binding sites in the monomeric enzyme, yellow bar in Fig. 1C) increases with valency. On the other hand, the solvent accessibility of the inhitopes (Fig. 2A) decreases with valency. In general, it can be expected that these two compensatory trends are expected to cancel out (to some extent). There are, however, some particularities. For example, glycoclusters **3** and **4** (both *n*=12) display rather similar normalized distributions of DNJ-DNJ distances. The accessibility of the DNJ heads is, however, very different (see Figure 2A-C). Cluster **3** features 12 *tripod* branches, each with two ghost side-arms, *i*.*e*., arms that are not decorated by an inhitope. These ghost side-arms shield the remaining inhitope-decorated arm from the solvent (Figure 2B). In glycocluster **4** with 12 *unipod* branches these ghost-arms are not present and the accessibility of the inhitopes is substantially increased with respect to cluster **3**. This could explain a higher chemical activity of the DNJ heads and therefore a higher inhibition potency of **4** with respect to **3**. Another particularity concerns the 12-valent glycoclusters **5a**-**c** that feature a *tripod* branch type with three inhitope-decorated arms. Obviously, inhitopes from the same branch cannot simultaneously bind at two different sites of the receptor (that are too far away). Or, in other words, inhitopes from the same branch (*tripod* case) are closer to each other than inhitopes from different branches. This is also seen by a shift of the normalized distribution of DNJ-DNJ distances to lower values (*i*.*e*., shift to the left in Figure 2A) when comparing **5c** with the *unipod* cluster **4**; the latter does not suffer from such geometric constraints. Thus, the degeneracy coefficient of clusters **5a**-**c** is in fact only 12x9=108 (due to geometric constraints) and therefore reduced with respect to **4** (12x11=132 without geometric constraints). This is reflected by a reduced inhibition potency. In principle, cluster **1** suffers also from these geometric constraints but the relative influence on the degeneracy coefficient is much less pronounced (36x33 with geometric constraints *vs* 36x35 without geometric constraints).

**Figure 2.**
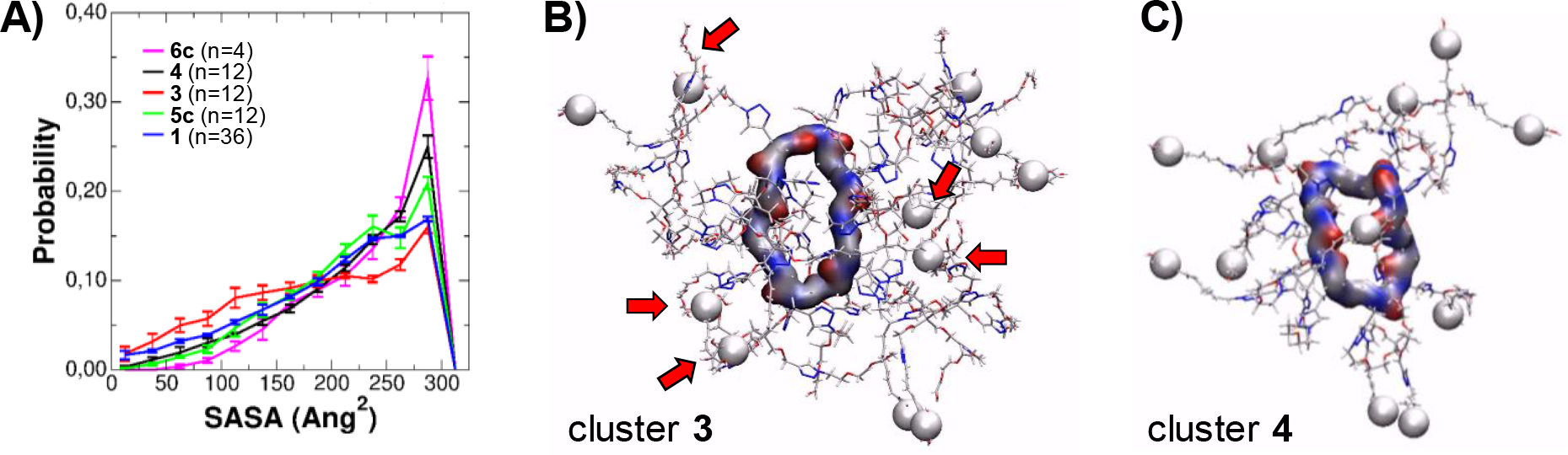
Solvent accessibility of DNJ inhitopes. A) Probability distribution of solvent accessible surface area (SASA) of the DNJ heads as calculated with a solvent probe radius of 2.4 Ang. B) Typical snapshot from the molecular dynamics simulation of cluster **3**. The backbone is shown as surface, the bonds of the branches as thin sticks and the nitrogen atoms of the DNJ heads as white balls. The arrows indicate inhitopes that are shielded from the side-arms. C) Same as subfigure B) but for cluster **4**.

### Limitations of the non-cooperative thermodynamic model

For the binding of the studied inhibitors to a single JBα-man molecule the radial topology assumption might be valid to some extent due to compensation effects. When comparing the experimental relative inhibition potencies with those calculated by Eq.7 (Fig. 3A), we realize, however, that the current thermodynamic model does not capture the full extent of the multivalent effect. In fact, the experimental *rp* values show an approximate *n*^3.5^-dependency for large values of *n* Eq. 7 provides only a *n*^2^-dependency. There are two potential explanations for this discrepancy:

**Figure 3:**
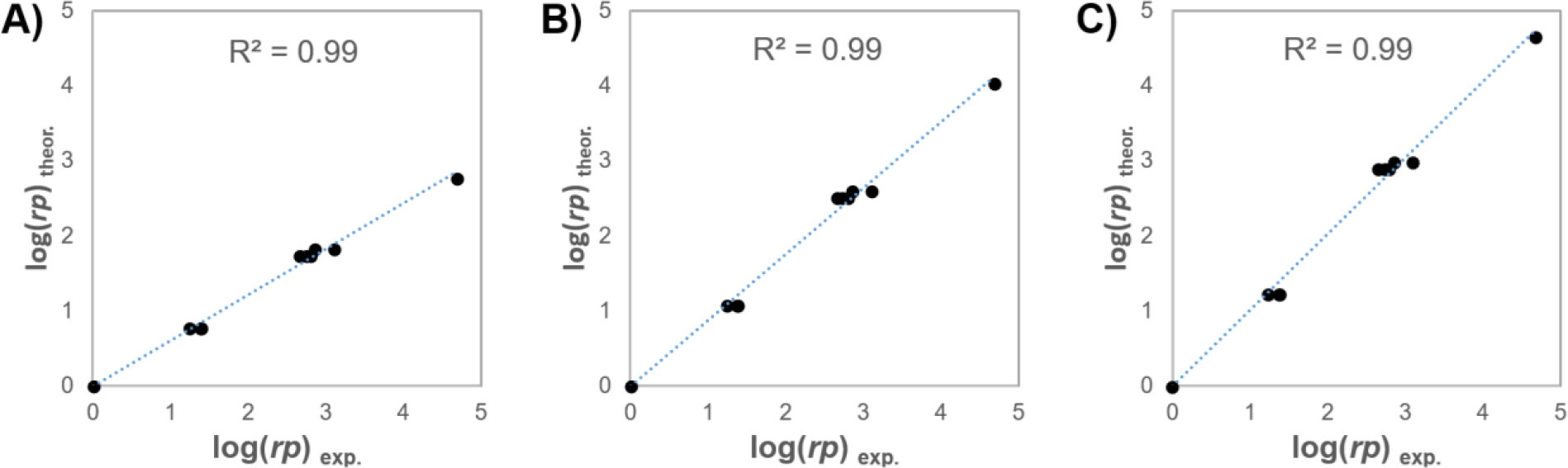
Correlation of experiment (abscissa) and theoretical model (ordinate) based on Eq.7 (A), Eq. 11 (B) and Eq. 10 with *p*=1.5 (C). For clusters **1** and **5a**-**c** we used a reduced degeneracy coefficient (see text). The main difference between these graphs is the increasing slope from 0.61 (A) to 0.88 (B) and 1.01 (C).

1. Possibility to form complexes with 2:1 stoichiometry (2 JBα-man receptors:1 multivalent inhibitor). These complexes feature four receptor binding sites which leads to higher order terms, *i*.*e*., additional *n*^3^- and *n*^4^-terms, to properly describe the *n*-dependency of the degeneracy coefficient.^12^ If 2:1 complexes were responsible for the *n*^3.5^-like dependency of *rp*, it would imply that the fraction of formed 2:1 complexes should increase with the valency. Indeed, high-valency clusters **1** and **4** are the only clusters of the series that have been found to form 2:1 complexes.^6^ On the other hand, both clusters feature a rather similar fraction of 2:1 complexes.^6^ A significantly larger fraction of 2:1 complexes would be expected for 36-valent cluster **1** in comparison to 12-valent cluster **4** (even if the geometric constraints of cluster **1** are taken into account). It should also be noted that cluster **4** is the only 12-valent cluster that forms a 2:1 complex. In the case of 12-valent cluster **3** the solvent accessibility and therefore the chemical activity of the DNJ heads is probably too low to form such a 2:1 complex. Note that a reduced chemical activity of the inhitope impacts the formation of a fully-inhibited 2:1 complex by the power of 4 (∼ *k*_on_^4^). In the case of the 12-valent clusters **5a**-**c** the degeneracy coefficient is significantly reduced due to geometric constraints (see above). The degeneracy coefficient for a fully-inhibited 2:1 complex for cluster **4** is 12x11x10x9=11880 while for clusters **5a**-**c** it is only 12x9x6x3=1944. Finally, the experimental conditions are different for the measurement of the inhibition constant and the fraction of 2:1 complexes. It can therefore not be concluded with certainty that the formation of complexes with 2:1 stoichiometry is responsible for the *n*^3.5^-like dependency of *rp*.
2. Increasing favorable electrostatic interactions with increasing inhitope valency. At pH=5.5 the *apo* JBα-man features an overall negative charge of about -6.6±0.2*e* as obtained from continuous constant-pH molecular dynamics simulations.^17^ The inhibitors, however, are charged positively. For example, **6c** (n=4) and **5c** (n=12) are charged +2.4±0.3*e* and +5.4±0.3*e*, respectively, due to the partial protonation of the DNJ inhitopes. Due to their opposite charge, the enzyme and the neoglycoclusters attract each other, and the attractive force increases with increasing valency. At pH=4, on the other hand, the charge of the enzyme is 9.4±0.1*e*. (The isoelectric point of this enzyme is at about pH=5). The charge of the clusters 6c and 5c is +3.2±0.2*e* and +8.1±0.3*e*, respectively. Thus, at pH=4 repulsive electrostatic interactions are expected between the positively charged enzyme and the positively charged inhibitor and this repulsion would increase with increasing valency, *i*.*e*., with increasing positive charge.

If these electrostatic interactions played indeed a major role for the multivalent effect, then lowering of the pH from 5.5 to 4 would display a stronger impact on the inhibition potency for high valencies than for low valencies. To test this hypothesis, inhibition constants of two representative compounds have been measured at pH 4.^6^ *K*_i_ of the tetravalent cluster **6c** increases by a factor of about 80 from 2.3 μM (pH 5.5) to 184 μM (pH 4). On the other hand, *K*_i_ of the 12-valent **5c** increases by a factor of more than 400 from 97 nM (pH 5.5) to 42 μM (pH 4). Indeed, the cluster with the higher valency displays the stronger decrease in inhibition potency. As a consequence we decided to develop an extension of the thermodynamic model that empirically includes electrostatic effects.

### Extension to charged inhitopes

It is well known that electrostatics play a crucial role for the associations of macromolecules (in particular for the parameter *K*_*1*_ of Eq. 7). Usually the binding affinity is linearly correlated to the product of the total charge of the two interacting molecules. It has been shown by MC simulations, however, that for oppositely charged macromolecules in polar solvents this linear correlation breaks for larger net charges.^18^ By keeping the charge of one macromolecule fixed at a certain negative value and increasing the positive charge of the other molecule, the binding affinity displayed a logarithmic-like behavior. We can therefore propose the following dependence of *K*_1_ on *n*:

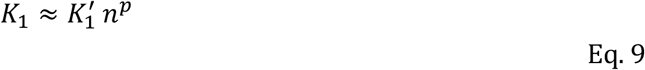

and we obtain:

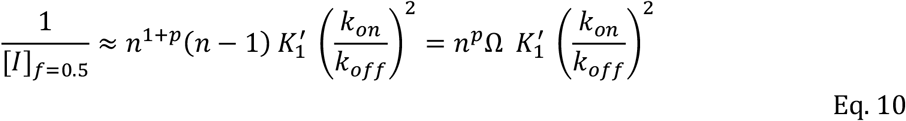

Where *K*_1_’ is the association constant of the encounter complex for a monovalent ligand and *p* is a parameter of the studied system (*i*.*e*., enzyme & type of inhitope). This parameter may also depend on the experimental conditions (*e*.*g*. pH, see below). With *p*=0 we recover Eq. 7. For *p*=1 we can derive the following simple expression for the relative inhibition potency:

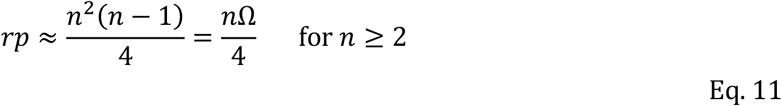

Here again the divalent cluster **7** was used as reference for the calculation of *rp*. Fig. 3B shows the correlation between Eq. 11 and the experimental *rp* values. Already this simple model provides an excellent description for the dependence of *rp* on *n*. A slightly better regression slope (=closer to unity) is obtained with *p*=1.5 (Fig. 3C)^6^ as used in the companion study.^6^

## Conclusion

Simplistic thermodynamic models can be applied to understand in parts the multivalent effect of JBα-man inhibition by neoglycoclusters. Microscopic structural differences such as the inhitope distributions and solvent accessibility can lead, however, to deviations from such simplistic models. An extension to the model has been proposed for charged inhitopes to take into account electrostatic contributions to the binding affinity. In summary, the studied multivalent inhibitors of this work and the herein developed extended thermodynamic model are unique in the sense that they open up a new dimension (electrostatics) in addition to the standard statistical (degeneracy) dimension. We anticipate that this approach can be applied to other protein targets and thereby pave the way to the design of new and more targeted multivalent inhibitors.

## Methods

### General

Simulations with implicit solvation model were carried out with the program CHARMM,^19^ version c45b2, on CPU Intel Xeon Gold 6126. Explicit-water simulations were performed with OpenMM,^20^ version 7.4, on GPU Nvidia RTX2080 with mixed precision. Continuous constant pH-simulations were carried out with the OpenMM-interface[ref] of CHARMM c45b2 on GPU Nvidia GTX1080. Input files for the enzyme (including two Zn^2+^ ions and four glycan modifications) were obtained from the CHARMM-GUI server^21^ with the PDB Reader module using the PDB entry 6B9P and the force field CHARMM36m. Parameters of the zinc (II) ions were those of Stote and Karplus.^22^ Topologies & parameters of the neoglycoclusters were also obtained from the CHARMM-GUI server using the Ligand Reader module (CGenFF^23^ parameters). All DNJ heads were constructed in their protonated state.

### Molecular dynamics simulation of neoglycoclusters

For each cluster we generated with CHARMM four different starting structures by simulated annealing using the implicit solvation model FACTS^24^ and its recommended simulation parameters. The initial model from CHARMM-GUI was first heated to 450 K within 1ns and then cooled down to 298.15K within 10ns using Langevin dynamics. This procedure was repeated four times with different initial velocities (ISEED) leading to four different starting structures for the subsequent simulations with OpenMM. Each structure was solvated in a cubic box of TIP3P water molecules (side length 5 nm) with periodic boundary conditions. Non-bonded interactions were cut off at 0.9 nm. Long range electrostatic interactions were treated by Particle-Mesh Ewald. Covalent hydrogen bonds were kept constrained. Equation of motion was integrated with OpenMM’s Langevin integrator at constant temperature (298.15 K). After a short equilibration of 10 ns in the NVT ensemble using a time step of 1 fs we performed 300 ns of MD in the CPT ensemble (1 atm) using a time step of 2 fs. The last 200 ns were used for analysis purposes, *i*.*e*., 100 ns served as equilibration. Each MD simulation (originating from a different starting structure) was first separately analyzed, and then mean values were obtained by averaging over the four separate analyses of each neoglycocluster. Error bars correspond to standard error of the mean.

### Constant pH-Simulations of neoglycoclusters and JBα-man enzyme

Similar to the explicit-water simulations we created different starting structures (5 in total) for the enzyme and the clusters **5c** and **6c** using the same simulated annealing protocol. Simulations were carried out, however, with a continuous constant pH method at pH-values of 4.0 and 5.5 using the GBSW solvation model with the approach of Brooks and coworkers.^25^ Standard settings were used for the GBSW model (*i*.*e*., ionic strength and surface tension coefficient were kept at zero). In the case of the enzyme a restraining force (CONS HARM) was applied to the non-hydrogen protein & zinc atoms during the simulated annealing runs (force constants of 10.0 and 1.0 kcal/mol/Ang^2^ for the backbone/Zn^2+^ and side-chains, respectively) to avoid unfolding of the enzyme. For the DNJ inhitope we used a pK_a_ of 6.94 as known from the literature (N-butyl-DNJ, miglustat).^9^ Following a calibration procedure,^26^ the continuous-pH parameters A and B were set to -60.3 and 0.453, respectively, for the DNJ inhitope. Parameter BARR was set to 1.75. For the amino-acid groups of the enzyme we used the standard pK_a_ values and continuous-pH parameters of the GBSW solvation model as provided in the CHARMM distribution. Each starting structure was then equilibrated for 5 ns at T=298.15K before a production run of 5 ns occurred. The restraining forces were reduced by a factor of 10 with respect to the simulated annealing simulations. For each simulation the mean charge of the cluster or the enzyme was obtained by averaging over all snasphots (saved every 500 ps). Then mean values and standard errors of the mean were obtained by averaging over the five simulations (corresponding to the five different starting structures).

## Acknowledgement

The computational part of this work was supported by research grants g2023a142c/g and g2022a262c/g from the Computer Center of the University of Strasbourg (CCUS). We are grateful to Prof. Jana Shen and Prof. Charles Brooks III for help on the continuous constant pH-MD code.

## Notes

### Competing Interest Statement

The authors have declared no competing interest.

## References

(1) Asano, N. Glycosidase Inhibitors: Update and Perspectives on Practical Use. Glycobiology 2003, 13 (10), 93R–104R. 10.1093/glycob/cwg090.

(2) Wadood, A.; Ghufran, M.; Khan, A.; Azam, S. S.; Jelani, M.; Uddin, R. Selective Glycosidase Inhibitors: A Patent Review (2012–Present). Int. J. Biol. Macromol. 2018, 111, 82–91. 10.1016/j.ijbiomac.2017.12.148.

(3) Compain, P. Multivalent Effect in Glycosidase Inhibition: The End of the Beginning. Chem. Rec. 2020, 20 (1), 10–22. 10.1002/tcr.201900004.

(4) Fasting, C.; Schalley, C. A.; Weber, M.; Seitz, O.; Hecht, S.; Koksch, B.; Dernedde, J.; Graf, C.; Knapp, E.-W.; Haag, R. Multivalency as a Chemical Organization and Action Principle. Angew. Chem. Int. Ed. 2012, 51 (42), 10472–10498. 10.1002/anie.201201114.

(5) Note 1: The reader is referred to the companion study (ref. 5) for more details on the exact chemical structure of the clusters, their synthesis and measurements of the inhibition constants against JBα-man.

(6) Liang, Y.; Schettini, R.; Kern, N.; Izzo, I.; Spichty, M.; Bodlenner, A.; Compain, P. Deconstructing Best-in-Class Neoglycoclusters as a Tool for Dissecting Key Multivalent Processes in Glycosidase Inhibition. Chem. – Eur. J. (submitted).

(7) Lepage, M. L.; Schneider, J. P.; Bodlenner, A.; Meli, A.; De Riccardis, F.; Schmitt, M.; Tarnus, C.; Nguyen-Huynh, N.-T.; Francois, Y.-N.; Leize-Wagner, E.; Birck, C.; Cousido-Siah, A.; Podjarny, A.; Izzo, I.; Compain, P. Iminosugar-Cyclopeptoid Conjugates Raise Multivalent Effect in Glycosidase Inhibition at Unprecedented High Levels. Chem. – Eur. J. 2016, 22 (15), 5151–5155. 10.1002/chem.201600338.

(8) Brumshtein, B.; Greenblatt, H. M.; Butters, T. D.; Shaaltiel, Y.; Aviezer, D.; Silman, I.; Futerman, A. H.; Sussman, J. L. Crystal Structures of Complexes of N-Butyl- and N-Nonyl-Deoxynojirimycin Bound to Acid β-Glucosidase. J. Biol. Chem. 2007, 282 (39), 29052–29058. 10.1074/jbc.M705005200.

(9) European Medicines Agancy. Assessment Report: Miglustat Dipharma, 2018. https://www.ema.europa.eu/en/documents/assessment-report/miglustat-dipharma-epar-public-assessment-report_en.pdf.

(10) Howard, E.; Cousido-Siah, A.; Lepage, M. L.; Schneider, J. P.; Bodlenner, A.; Mitschler, A.; Meli, A.; Izzo, I.; Alvarez, H. A.; Podjarny, A.; Compain, P. Structural Basis of Outstanding Multivalent Effects in Jack Bean α-Mannosidase Inhibition. Angew. Chem. Int. Ed. 2018, 57 (27), 8002–8006. 10.1002/anie.201801202.

(11) Kane, R. S. Thermodynamics of Multivalent Interactions: Influence of the Linker. Langmuir 2010, 26 (11), 8636–8640. 10.1021/la9047193.

(12) Kitov, P. I.; Bundle, D. R. On the Nature of the Multivalency Effect: A Thermodynamic Model. J. Am. Chem. Soc. 2003, 125 (52), 16271–16284. 10.1021/ja038223n.

(13) Huskens, J.; Mulder, A.; Auletta, T.; Nijhuis, C. A.; Ludden, M. J. W.; Reinhoudt, D. N. A Model for Describing the Thermodynamics of Multivalent Host–Guest Interactions at Interfaces. J. Am. Chem. Soc. 2004, 126 (21), 6784–6797. 10.1021/ja049085k.

(14) Sohrabi-Jahromi, S.; Söding, J. Thermodynamic Modeling Reveals Widespread Multivalent Binding by RNA-Binding Proteins. Bioinformatics 2021, 37 (Supplement_1), i308–i316. 10.1093/bioinformatics/btab300.

(15) von der Lieth, C.-W.; Frank, M.; Lindhorst, T. K. Molecular Dynamics Simulations of Glycoclusters and Glycodendrimers. Rev. Mol. Biotechnol. 2002, 90 (3), 311–337. 10.1016/S1389-0352(01)00072-1.

(16) Perez, S.; Makshakova, O. Multifaceted Computational Modeling in Glycoscience. Chem. Rev. 2022, 122 (20), 15914–15970. 10.1021/acs.chemrev.2c00060.

(17) Note 2: These simulations were carried out at low ionic strength which explains the somehow low absolute charge of the enzyme.

(18) Jönsson, B.; Lund, M.; Barroso da Silva, F. L. Electrostatics in Macromolecular Solutions. In Food Colloids: Self-Assembly and Material Science; RSCPublishing, 2007.

(19) Brooks, B. R.; Brooks III, C. L.; Mackerell, A. D.; Nilsson, L.; Petrella, R. J.; Roux, B.; Won, Y.; Archontis, G.; Bartels, C.; Boresch, S.; Karplus, M.; others. CHARMM: The Biomolecular Simulation Program. J Comput Chem 2009, 30 (10), 1545–1614. 10.1002/jcc.21287.

(20) Eastman, P.; Swails, J.; Chodera, J. D.; McGibbon, R. T.; Zhao, Y.; Beauchamp, K. A.; Wang, L.-P.; Simmonett, A. C.; Harrigan, M. P.; Stern, C. D.; Wiewiora, R. P.; Brooks, B. R.; Pande, V. S. OpenMM 7: Rapid Development of High Performance Algorithms for Molecular Dynamics. PLOS Comput. Biol. 2017, 13 (7), e1005659. 10.1371/journal.pcbi.1005659.

(21) Jo, S.; Kim, T.; Iyer, V. G.; Im, W. CHARMM-GUI: A Web-Based Graphical User Interface for CHARMM. J. Comput. Chem. 2008, 29 (11), 1859–1865. 10.1002/jcc.20945.

(22) Stote, R. H.; Karplus, M. Zinc-Binding in Proteins and Solution - a Simple but Accurate Nonbonded Representation. Proteins 1995, 23, 12–31. 10.1002/prot.340230104.

(23) Vanommeslaeghe, K.; Hatcher, E.; Acharya, C.; Kundu, S.; Zhong, S.; Shim, J.; Darian, E.; Guvench, O.; Lopes, P.; Vorobyov, I.; MacKerell, A. D. CHARMM General Force Field (CGenFF): A Force Field for Drug-like Molecules Compatible with the CHARMM All-Atom Additive Biological Force Fields. J. Comput. Chem. 2010, 31 (4), 671–690. 10.1002/jcc.21367.

(24) Haberthuer, U.; Caflisch, A. FACTS: Fast Analytical Continuum Treatment of Solvation. J Comput Chem 2008, 29, 701–715.

(25) Lee, M. S.; Salsbury, F. R.; Brooks, C. L. Constant-PH Molecular Dynamics Using Continuous Titration Coordinates. Proteins 2004, 56 (4), 738–752. 10.1002/prot.20128.

(26) Henderson, J. A.; Liu, R.; Harris, J. A.; Huang, Y.; Oliveira, V. M. de; Shen, J. A Guide to the Continuous Constant PH Molecular Dynamics Methods in Amber and CHARMM [Article v1.0]. Living J. Comput. Mol. Sci. 2022, 4 (1), 1563–1563. 10.33011/livecoms.4.1.1563.

